# qFit 3: Protein and ligand multiconformer modeling for X-ray crystallographic and single-particle cryo-EM density maps

**DOI:** 10.1101/2020.09.03.280222

**Authors:** Blake T. Riley, Stephanie A. Wankowicz, Saulo H. P. de Oliveira, Gydo C. P. van Zundert, Daniel Hogan, James S. Fraser, Daniel A. Keedy, Henry van den Bedem

## Abstract

New X-ray crystallography and cryo-electron microscopy (cryo-EM) approaches yield vast amounts of structural data from dynamic proteins and their complexes. Modeling the full conformational ensemble can provide important biological insights, but identifying and modeling an internally consistent set of alternate conformations remains a formidable challenge. qFit efficiently automates this process by generating a parsimonious multiconformer model. We refactored qFit from a distributed application into software that runs efficiently on a small server, desktop, or laptop. We describe the new qFit 3 software and provide some examples. qFit 3 is open-source under the MIT license, and is available at https://github.com/ExcitedStates/qfit-3.0.

## Introduction

Conformational dynamics play an essential role in many aspects of protein function, including ligand binding, allostery, and enzyme turnover ^1,2^. In each of these processes, the protein does not adopt a single conformation, but rather a conformational ensemble including a number of low-energy states. This ensemble can then be redistributed or reshaped by small-molecule binding, post-translational modifications, or other perturbations, thereby controlling biological function. To fully understand the fundamental interplay between protein conformational heterogeneity and function, it is necessary to develop experimental and computational techniques to reveal alternative protein conformations in atomic detail.

X-ray crystallography is a powerful tool for addressing this need. Because individual protein molecules in the crystal lattice sample different conformations, there is a growing appreciation that crystallographic electron density maps contain a wealth of information about sparsely populated, alternative protein conformations ^3^. Moreover, crystallography is undergoing an experimental renaissance: new tools are emerging with the potential to bias conformational distributions in crystals and gain new mechanistic insights into the links between protein dynamics and function.

For example, crystallographic datasets collected across multiple temperatures — as opposed to at a single cryogenic temperature — often reveal ensembles with more conformational diversity ^4–8^, including at dynamic enzyme active sites ^9^. High-throughput crystallographic protein:ligand screening can identify otherwise undetectable low-occupancy ligand-bound protein states ^7,10,11^. And time-resolved diffraction experiments, triggered by a variety of stimuli ^12–15^, can offer detailed windows into how protein conformational ensembles evolve in real time. Time-resolved experiments are becoming more accessible as serial microcrystallography experiments can take place not only at ray free-electron lasers, but also at third-generation synchrotrons with microfocus beamlines ^16^. Serial microcrystallography can also help reveal alternative protein states by dissecting distinct crystal polymorphs within the microcrystal population ^17^. These advances, coupled with an ever-growing level of automation and faster X-ray detectors ^18^, are yielding larger amounts of data that highlight the need for automated (rather than manual) computational methods for modeling alternative conformations and their correlations in electron density maps.

In parallel to the renaissance for X-ray crystallography, cryo-electron microscopy (cryo-EM) is in the midst of a “resolution revolution” ^19^. Recently, cryo-EM structures of two different systems at “atomic resolution” (1.2–1.25 Å) ^20,21^ demonstrated how far this method has come in recent years. Similar to electron density maps from X-ray crystallography, Coulomb potential maps from cryo-EM reveal evidence for alternative protein states, which in this case are sampled by individual protein molecules on the microscopy grid. Unfortunately, so far no methods exist for unbiased and automatic modeling of correlated alternative conformations in cryo-EM maps. Additionally, many cryo-EM structures feature large protein complexes with thousands of amino acids, posing a significant challenge to traditional model building approaches. Efficient, automated algorithms ^22^ could meet this challenge for cryo-EM.

There is thus a clear need for computational model-building methods that better explain X-ray and cryo-EM data by incorporating alternative conformations. Protein conformational heterogeneity can be represented using various approaches, including B-factors, multi-copy ensembles, or multiconformer models ^1^. First, B-factors are present for every atom in the Protein Data Bank (PDB) ^23^ file format. Theoretically, B-factors represent the harmonic, thermal displacement of each atom about its mean position, either isotropically with one parameter or anisotropically with six parameters ^24^. However, in practice, B-factors often absorb uncertainty in a more general sense about each atom’s position, and are insufficient representations of anharmonic motions such as transitions between side-chain rotamers ^25^. Second, multi-copy ensemble models consist of some number (>1) of full, independent copies of the protein with distinct xyz coordinates and B-factors that collectively explain the experimental data ^26^. Ensemble models can successfully describe discrete conformational heterogeneity such as rotamer transitions -- but they unnecessarily inflate the number of model parameters for those regions of the protein with essentially a single, unique conformation ^27^. Finally, multiconformer models lie somewhere in the middle in terms of model complexity. A multiconformer model represents local, anharmonic features in the data with a small number (2–5) of discrete conformations, but represents regions of the protein that show little to no evidence of flexibility with a single conformation. These conformations are assigned labels (“alternative locations” or “altlocs”), such as A, B, etc., with corresponding occupancies in the PDB format on a per-atom basis. Groups of atoms whose alternate positions are correlated (side chains, stretches of contiguous backbone, collective exchange across an active site, etc.) are assigned the same label and occupancy. When constructed in a parsimonious manner, multiconformer models can limit a model’s complexity while maximizing its explanatory power.

To efficiently generate parsimonious multiconformer models for protein X-ray crystal structures, we previously introduced the software package qFit ^28^. Besides providing mechanistic insights, for example by revealing hidden protein contact signaling networks ^29^ and allosteric pathways ^7^, multiconformer qFit models have also established that the conformational ensemble at room temperature is not dominated by radiation damage ^30^, and that the effect of crystal dehydration on the conformational ensemble is similar to that of cryocooling ^31^. We recently introduced multiconformer treatment of ligands in complex with proteins in a standalone version, *qFit-ligand* ^32^. However, previous versions of qFit were computationally demanding (requiring a high-performance computing cluster), and were restricted to density maps from X-ray crystallography only, among other limitations.

Here we report a new, refactored version of qFit, which we call qFit 3, with several key improvements. qFit 3 operates on maps from either X-ray crystallography or cryo-EM. It combines multiconformer modeling of proteins and of ligands complexed with proteins (from *qFit-ligand*) in a single software package written in Python. The software distribution includes a script to refine the multiconformer model generated by qFit with Phenix ^33^. Importantly, we reduced the runtime by two orders of magnitude. qFit 3 typically runs for a ∼300 residue protein in several hours on a laptop, making it significantly more accessible to users.

Overall, qFit 3 reveals hidden alternative conformations in protein structures in a rapid, automated, and unbiased manner. This new software will allow a broader array of users to explore conformational heterogeneity in their systems of interest. It will also smooth the path toward integrating new and exciting types of structural biology data, including series of datasets related by temperature, ligands, or time, as well as biologically important and/or large protein systems from X-ray free electron lasers (XFELs) or cryo-EM. qFit 3 will thus empower novel studies of the relationship between protein dynamics and biological function.

## Results

qFit was completely refactored in the Python programming language and released as open-source software; see Methods and the GitHub repository (https://github.com/ExcitedStates/qfit-3.0) for more details. A typical qFit 3 workflow is illustrated in **Figure 1**. qFit 3 takes as minimal input a starting model and either a real-space map in the MRC/CCP4 format or map coefficients in the MTZ format. For X-ray crystallography, the preferred map is a composite omit map to minimize model bias, which can be readily generated with Phenix. For cryo-EM, the input is a real-space map together with the resolution of the data and a flag to use electron scattering factors for generating synthetic densities. qFit 3 relies on a sample-and-select procedure based on constrained optimization to identify alternative conformations of proteins and their ligands. To ensure optimal model selection and prevent overfitting, qFit 3 evaluates increasing model complexities, selecting the model with the lowest Bayesian Information Criterion (BIC). qFit 3 now also provides all functionality to model ligand alternate conformations, previously available separately in *qFit-ligand*. A distinctly important new feature is qFit 3’s capability to model alternate conformations into cryo-EM maps. Numerous additional options and details are described in the Methods section and can be found in the qFit 3 GitHub repository. Here, we demonstrate typical use cases of qFit for protein systems and their ligands. All analyses in this section used default parameters, unless otherwise stated.

**Figure 1:**
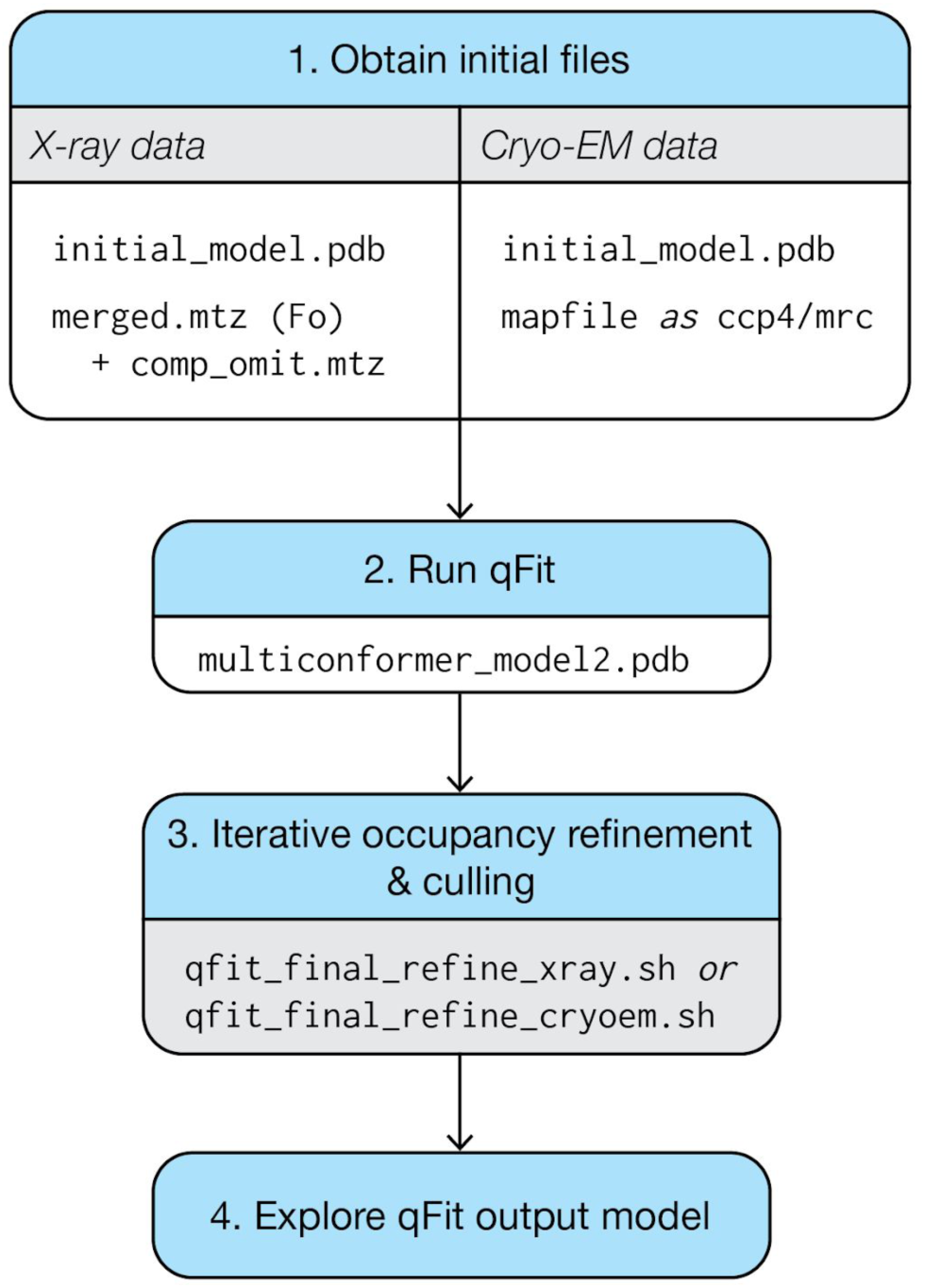
Usage flowchart for qFit 3 for either protein or ligand inputs and for either X-ray or cryo-EM data. (1) qFit requires an initial model and map information. In the case of X-ray diffraction data, qFit will require both the structure factors and a high-quality, unbiased map, such as a composite omit map. (2) With these files, qFit will generate a parsimonious model (multiconformer_model2.pdb) containing the fewest number of sampled conformers that explain the experimental data. (3) This intermediate/preliminary model should proceed through an iterative procedure to refine the occupancies of conformers in the model, and cull those conformers that have <9% occupancy. (4) The resulting model can then be used to explore conformational diversity. See **Supplementary Figures 1 & 2** for more detail on usage for X-ray vs. cryo-EM data.

We first carried out qFit 3 modeling on a previously deposited cryogenic X-ray structure of a protein tyrosine phosphatase, PTPN18 (PDB ID: 2oc3) ^34^. While the deposited model includes ten residues with alternate conformers, a difference density map shows unmodeled positive density over 3σ around Phe30 and Gln34 **(Figure 2A, left panel)**. qFit 3 models suggest that an alternate conformer for Phe30 and an ensemble of three side-chain conformers for Gln34 better fit the density, and reduce nearby difference density peaks (**Figure 2A, right panel)**. Running on a quad-core processor, qFit sampled and selected alternative conformations for this 290-residue protein in 12.75 hours.

**Figure 2:**
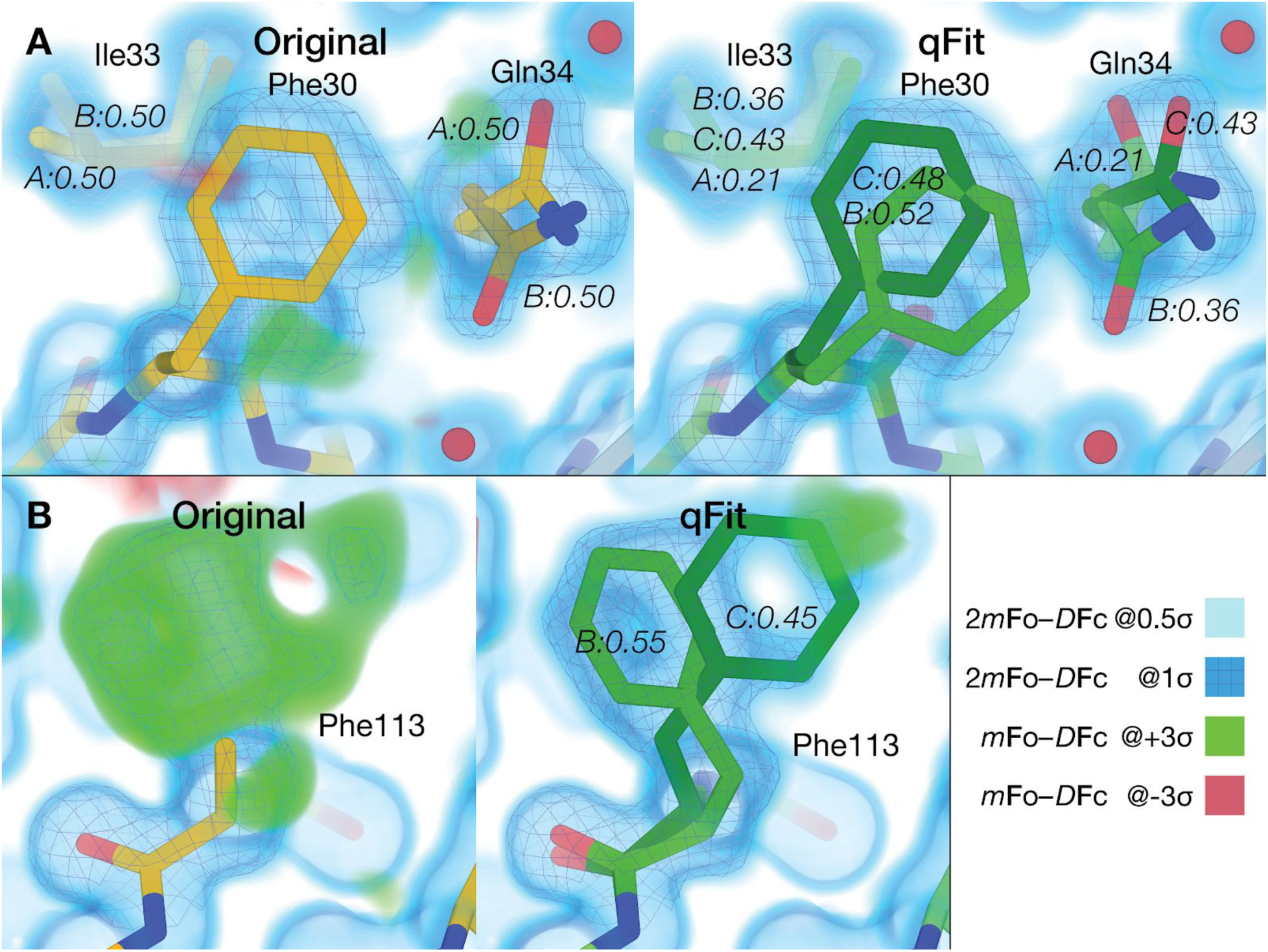
qFit 3 recapitulates deposited alternate conformations in X-ray crystallography density maps, and suggests additional conformations to explain unmodeled density. A). *Left*: PTPN18 (PDB ID: 2oc3) displays regions of unmodeled density near Phe30 and Gln34 in the deposited *m*Fo-*D*Fc difference density map at +3σ (green cloud). These are visible in a 2*m*Fo-*D*Fc composite omit density map contoured at 1σ (blue mesh), which is clarified by a low-density 0.5σ contour (blue cloud). *Right*: qFit 3 adds extra conformers to model these residues. Gln34 is modeled by three conformers (corresponding to the rotamers **mm**110, **mt**0, **mt**0 ^25^); Phe30 is also modeled by two conformers (both in the “favored” **t**80 rotamer space). The distance between Phe30 and Gln34 doesn’t lead to steric hindrance between any of the conformers of either residue. Note that qFit 3 sets the minimum number of conformers in Ile33 to three (because of Gln34) to ensure backbone consistency; Phe30 is part of another backbone segment. B) *Left*: Following the methodology in qFit 2 ^35^, Phe113 was truncated at Cβ and refined. Both the composite omit map and the difference map indicated the presence of at least two conformers for this residue. *Right:* qFit 3 sampled and selected two conformers of Phe113 (matching the two known ones) to explain the density in the composite omit map.

The default algorithm of qFit 3 changed slightly compared to earlier versions. Previously, each amino acid in turn was truncated at the Cβ atom and refined anisotropically. This had two advantages: 1) it generally positioned the Cβ atom at the peak *average* density of potential alternate conformations, and 2) the anisotropy of the atomic displacement parameter provided guidance for backbone motions. Although this earlier version often better captured subtle backbone movements, it led to significant increased computational expense and complexity ^35^. Nonetheless, the present version of qFit can be made to mimic the behavior of the earlier algorithm on a single residue by providing an alternative input. A thoroughly-tested room-temperature structure of the peptidyl-prolyl cis-trans isomerase CypA (PDB ID: 3k0n) displays multiple conformers for Phe113 ^9^. Starting from a single conformer (**Figure 2B, left panel**), we truncated Phe113 at Cβ, refined the structure anisotropically, calculated a composite omit map, and used this as input to qFit 3. This pre-processing enabled qFit to recapitulate the alternative conformations observed in the published model (**Figure 2B, right panel**). With Phe113 in place, qFit 3 ran in 460 min over the other 161 residues. This computationally expensive pre-processing procedure is provided as an option, and improved backbone modeling will be a focus of future development (Discussion).

qFit 3, for the first time, also accepts cryo-EM density maps as input. We have adopted the simplified scattering factor calculation of averaging the contributions of all atoms to calculate synthetic maps, as is used in real-space refinement in Phenix ^36^. As an example application of this new functionality, we ran qFit 3 on two ultra-high-resolution cryo-EM structures: β3 GABA receptor ^21^ (1.2 Å resolution) and apoferritin ^20^ (1.7 Å resolution). qFit 3 was run on both chain A and the entire structure for both examples. Chain A of apoferritin (176 residues) had a runtime of 112 minutes using four cores.

For these examples, qFit 3 captured both previously modeled and newly modeled alternative conformations (**Figure 3)**. Within chain A, there were originally 19 residues with modeled alternative conformers. qFit 3 successfully identified alternate conformations for 16 (84.2%) of these residues and suggested 66 additional residues with alternative conformations. In **Figure 3A**, we demonstrate the ability of qFit 3 to recapitulate alternative conformers in Ser124. In **Figure 3B**, we demonstrate the ability of qFit 3 to detect a new alternative rotamer for Gln14 (**pt0** and **mm-40** ^25^, **RMSF 1**.**16 Å)**.

**Figure 3:**
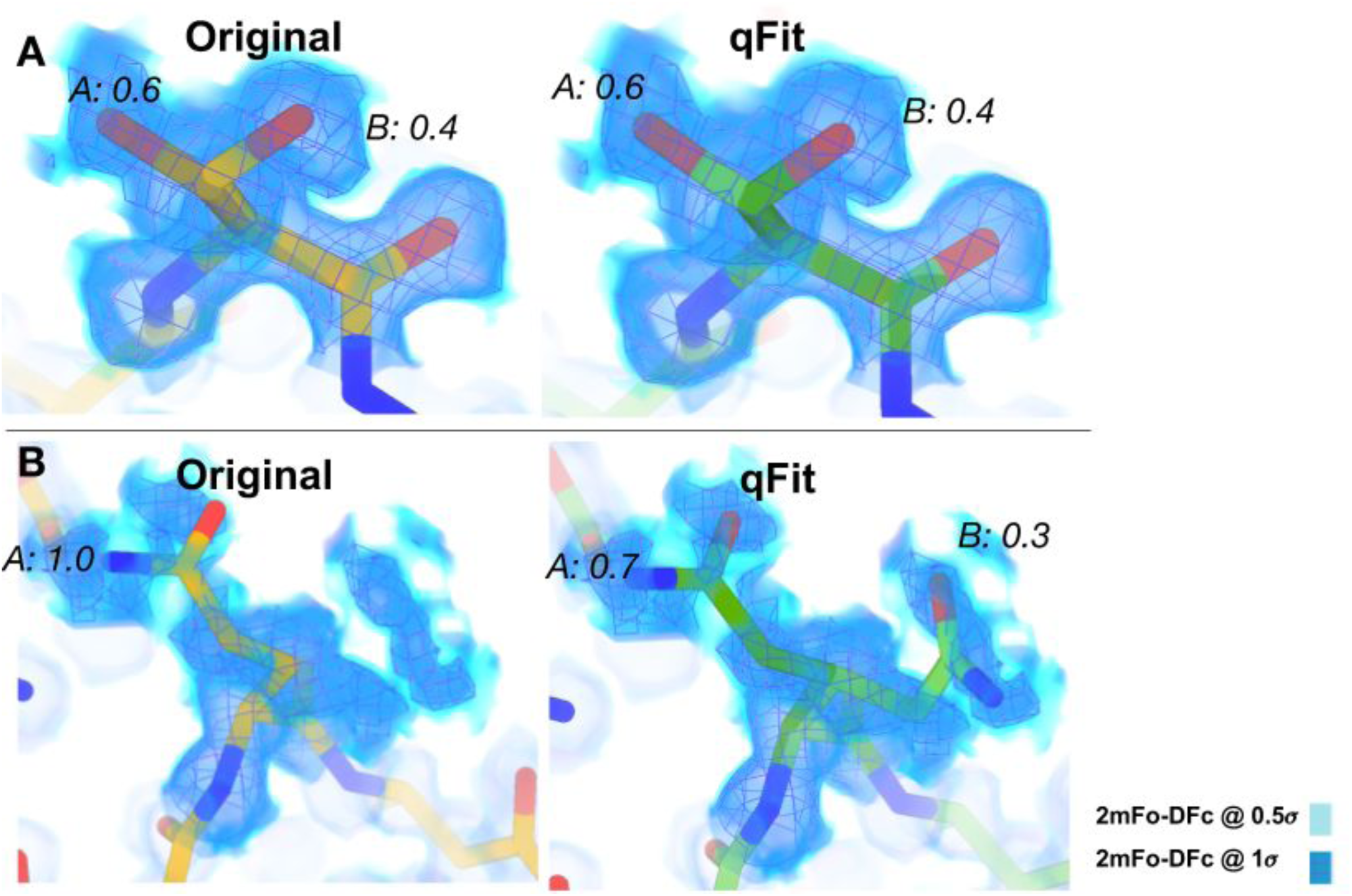
qFit 3 recapitulates deposited alternate conformations in cryo-EM density maps, and suggests alternate conformations to explain noisy data. A) *Left*: Deposited alternative conformations for Ser113 in a high-resolution published cryo-EM structure of apoferritin (PDB ID: 6v21). These are visible in a 2*m*Fo-*D*Fc composite omit density map contoured at 1σ (dark blue cloud) and at 0.5σ (light blue cloud and blue mesh). *Right*: qFit 3 and subsequent refinement successfully modeled identical alternative conformations. Occupancies are indicated in italics. B) *Left*: Deposited single conformation for Gln14 in the same structure of apoferritin. *Right*: qFit 3 and subsequent refinement identifies the original conformer, plus an alternative conformer (**mt** and **tt** rotamers ^25^).

Additionally, qFit 3 can determine alternative conformations of ligands ^32^. Distinct ligand conformations can play an important role in determining binding affinities, activity, and disassociation from the protein. Visualizing ligand alternate conformations can help determine the role of entropy in binding affinity, or help guide lead optimization in drug discovery ^37^. *qFit-ligand* takes a model, map, and information about the position of the ligand of interest (chain and residue number). The output is a set of conformations of the ligand. In **Figure 4**, we show two examples of ligands taking on multiple conformations to two different proteins, CDK2^38^ and Human Leukotriene A4 Hydrolase^39^.

**Figure 4:**
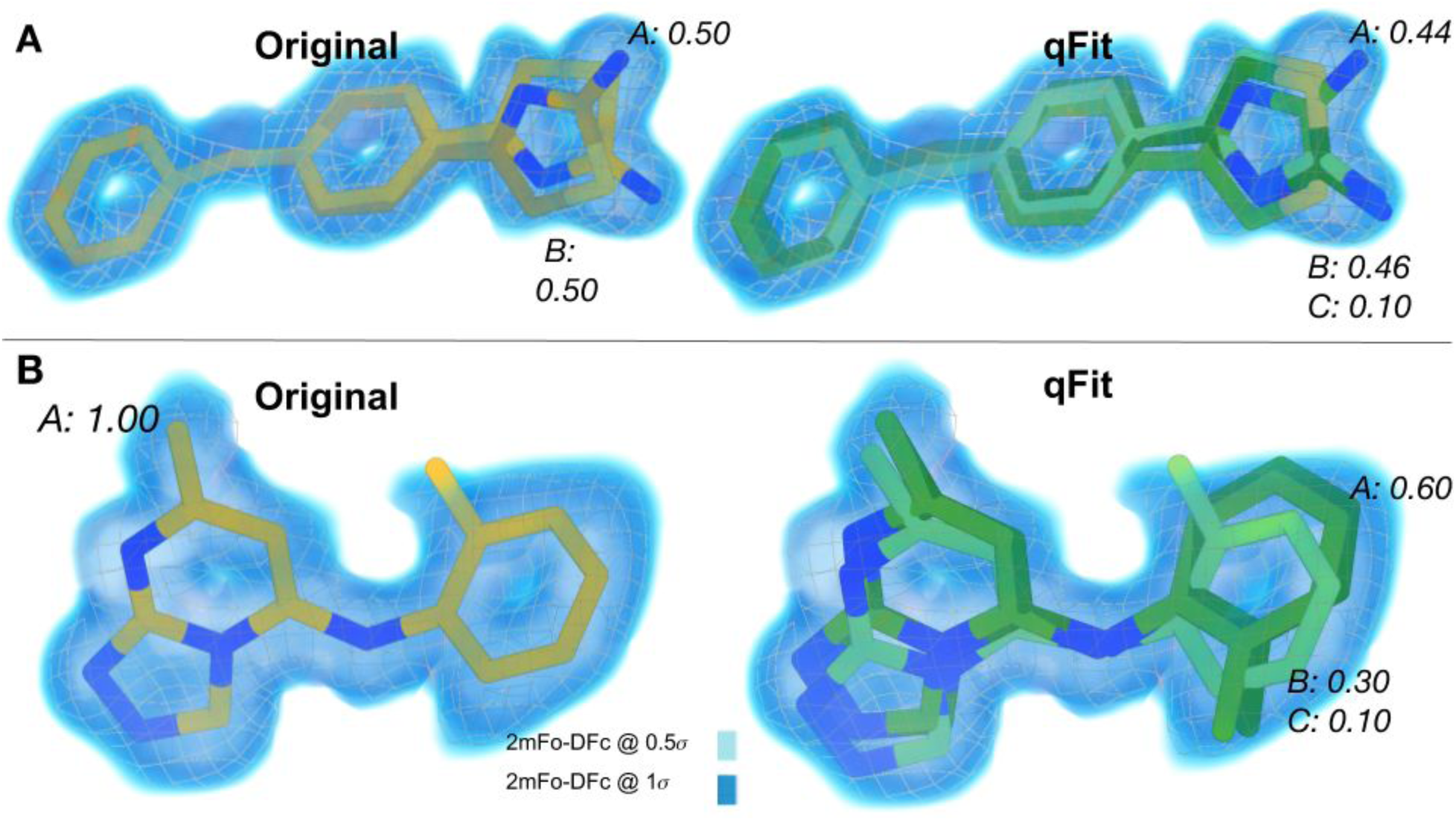
qFit 3 generates occupancy-weighted multiconformer models for bound ligands. A) *Left:* Deposited alternative conformations of thiazolylpyrimidine, an inhibitor of CDK2, in a co-crystal structure (PDB ID: 5hq5). The 2*m*Fo-*D*Fc composite omit density map is contoured at 1σ (dark blue cloud) and at 0.5σ (light blue cloud and grey mesh). Occupancies of alternative conformations are labeled in italics. *Right: qFit-ligand* successfully identifies both deposited alternative thiazolylpyrimidine conformations, as well as an additional, similar conformer. B) *Left:* Deposited conformation of 4-(4-benzylphenyl)thiazol-2-amine, an epoxide hydrolase selective inhibitor, co-crystallized with human Leukotriene A4 Hydrolase (PDB ID: 4l2l)^39^. *Right: qFit-ligand* models both the deposited 4-(4-benzylphenyl)thiazol-2-amine conformation and suggests two additional conformations that, unlike the deposited conformation, fit entirely within the 1σ density contour.

## Discussion

qFit 3 is a significantly faster implementation of the qFit algorithm that can now run on commodity computer hardware like a laptop. It is open-source and freely available, with simple installation instructions. qFit 3’s speed enables application of the qFit approach to series of multiple datasets generated by new high-throughput methods in crystallography; to large, increasingly high-resolution cryo-EM structures with many thousands of amino acids; and to many more structural bioinformatics studies that focus on conformational heterogeneity.

Although qFit 3 can be run in an automated fashion on large (numbers of) structures, the user should apply caution in interpreting its multiconformer models. False positives can occur when qFit 3 selects spurious alternative protein conformations based on density that corresponds to other atoms such as water molecules. False negatives can occur when qFit 3 fails to sample backbone conformational space sufficiently. Development of qFit is ongoing and the user community is invited to contribute to the open-source project at https://github.com/ExcitedStates/qfit-3.0.

To improve qFit further, we envision several new developments. For example, qFit’s backbone sampling methodology has ample room for improvement. Currently in qFit, each amino acid’s backbone is translated along the principal axes of the anisotropic ellipsoid of the Cß atom (or O for Gly), while closure of the backbone is maintained by torsion-based nullspace inverse kinematics, thus positioning it to accommodate suitable alternative side-chain rotamers (Methods). Although this current backbone sampling is powerful for capturing small-scale motions, it is limited in its ability to capture larger ones (**Figure 2B**). A suite of backbone sampling methods in qFit, ranging from backrubs ^40^ and helix “shear” ^41,42^ to inverse-kinematics-based loop modeling ^43^, would be able to overcome this limitation. These new methods will allow qFit to model alternative conformations that are related to each other by larger, biologically relevant motions, as with loops in protein tyrosine phosphatase 1B (PTP1B) ^7^ and helices in isocyanide hydratase (ICH) ^15^. A related challenge is that hierarchical alternative conformations — such as alternative loop or helix backbone positions that each have alternative side-chain rotamers — are not supported in the existing PDB format. It may be possible to use additional restraints to bypass this limitation, as with refinement of the multi-state models from PanDDA ^44^, which are conceptually related but distinct from the multiconformer models from qFit. Alternatively, the new PDBx/mmCIF format that was recently adopted by the PDB could be used to explicitly define hierarchical relationships between alternative conformations.

Another important direction is improving ligand models, and correlating protein alternate conformations with alternate ligand binding modes. Currently, qFit lacks chemical knowledge of ligand atoms such as hybridisation and protonation. Incorporating this knowledge, for example with the help of sophisticated force fields that work in tandem with crystallography maps ^45^, will greatly improve ligand model quality and help determine the precise interactions between protein and ligand.

Finally, the problem of compositional heterogeneity must be addressed. Some of the alternative conformations in the protein may be in response to the ordering of other components in the unit cell (heteroatoms such as ligands, crystallographic additives, and solvent). While multi-dataset approaches, such as PanDDA ^11^, may increase confidence in modeling partially occupied ligands and crystal additives, addressing the problem of partially occupied solvent may be bootstrapped by using stereotypical interactions in a solvated rotamer library ^46^. Solving this problem will also help to better define the border between proteins or ligands and bulk solvent ^47^, which is likely to be key to reducing the “R-factor gap in crystallography” ^48^.

## Conclusion

X-ray crystallography and cryo-electron microscopy remain the dominant experimental techniques to obtain structural information for proteins and their complexes with other macromolecules or with ligands, like therapeutic chemical compounds. New, emerging experimental techniques in X-ray crystallography and ever-increasing resolution limits in cryo-EM can reveal an ensemble of protein and ligand conformations that can provide insights into molecular mechanisms and function. qFit 3 automates interpreting an ensemble from X-ray or cryo-EM density maps, and generates an unbiased, internally consistent, parsimonious model of conformational heterogeneity. We refactored qFit with a specific focus on efficiency and ease-of-use, so that it effortlessly installs and runs on a standard laptop to facilitate advanced interpretation of experimental structural biology data.

## Methods

### qFit algorithm

#### Overview

qFit samples a large number of conformers and uses a deterministic approach to select a small ensemble of these conformers that parsimoniously explains local density. The method starts from an initial single-conformer model and generates candidate conformers for each residue/ligand in the initial structure. It evaluates all possible combinations of these conformers to determine an optimal ensemble. A final relabeling step ensures that conformers of different residues/ligands have consistent altloc labels. For all analyses in this manuscript, default parameters were used unless otherwise stated. **Figure 5** provides a graphical overview of both the *qFit-protein* and *qFit-ligand* algorithms, the two main command-line utilities of the qFit 3 package for automatic multiconformer modeling of proteins or ligands.

**Figure 5:**
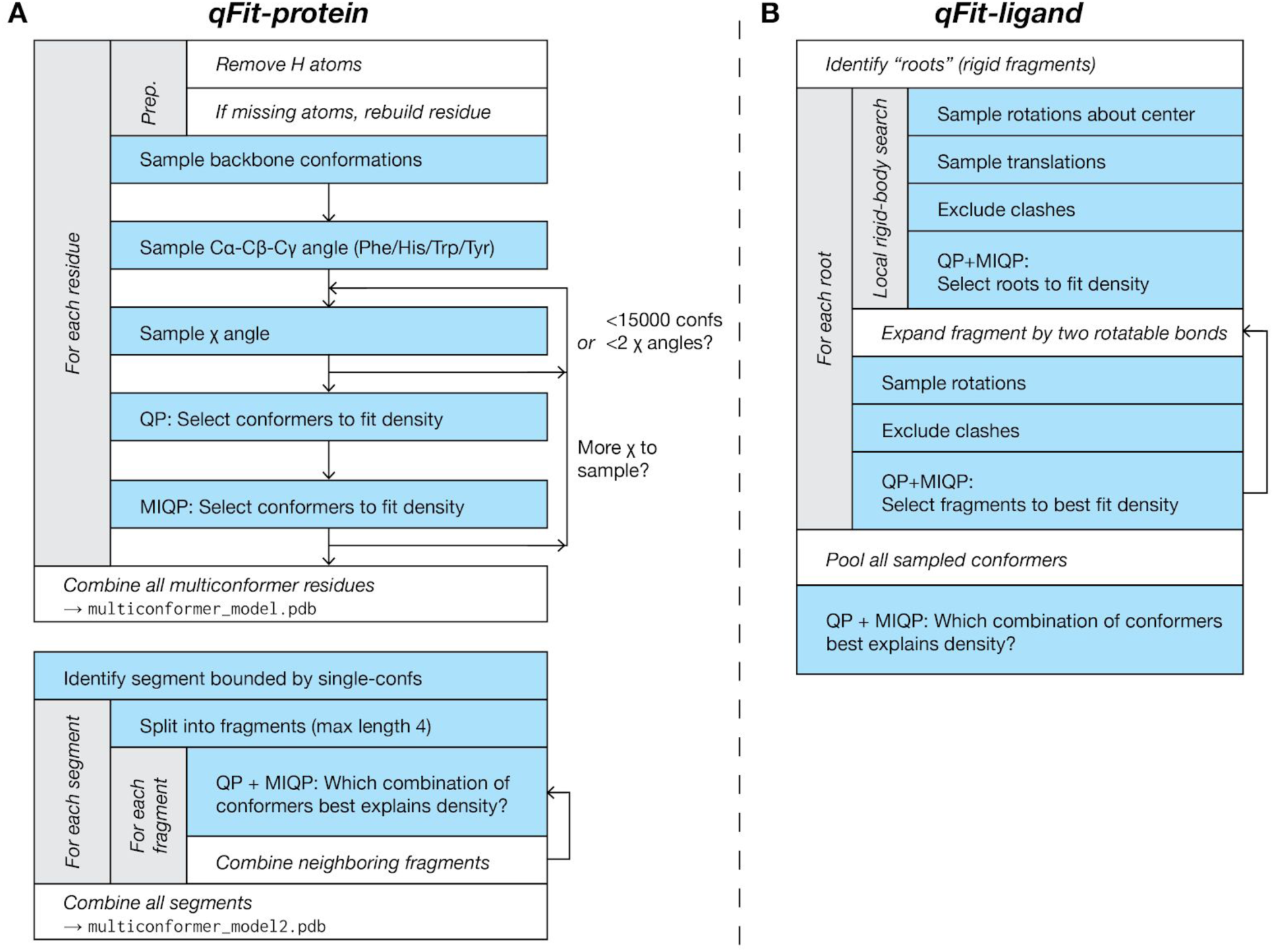
A flowchart of the sample-and-select protocols for (A) *qFit-protein*, and (B) *qFit-ligand*. See **Supplementary Figure 3** for a flowchart of the subsequent refinement stage.

#### Input

The qFit 3 protocol accepts input density maps or map coefficients in several commonly accepted crystallographic or cryo-EM file formats (MTZ, CCP4). For best performance, we recommend the use of a composite omit map for crystallographic densities ^49^. All runs of qFit 3 on crystal structures described in this manuscript used an input composite omit map generated with the *phenix*.*composite_omit_map* command from the Phenix software suite ^33^. Refinement was carried out on each partial model (*omit-type=refine*) and default parameters were used for this calculation. qFit 3 also expects a PDB file containing the structure of interest as input. Hydrogens are automatically removed to provide uniform treatment of input models. Note that during the final refinement stage, hydrogens will be (re-)added (see Final refinement script). For analyses described in this manuscript, we removed all alternate conformers (except for altloc A) using the *phenix*.*pdbtools* executable and used the resulting single-conformer input structure as input for all subsequent modeling.

#### Map treatment

qFit 3 converts the input maps to absolute scale following the protocol described in reference ^50^. The software creates a lookup table corresponding to the theoretical spatial density value distribution for each atomic element for radial shells spaced at 0.01 Å. The mask radius for this calculation is resolution-dependent (default radius = 0.5 Å + resolution/3). qFit 3 indirectly avoids clashes during sampling by means of a real-space density subtraction. It uses all atoms whose conformations are not being sampled to calculate a density map to perform this real-space subtraction. This prevents undesirable modeling into density from neighboring residues/side chains.

The mask radius and an option to use excluded volume for clash detection instead, as detailed in ^32^, can be determined via the command line. Different sets of scattering factors are used for electron density maps from X-ray crystallography vs. Coulomb potential maps from cryo-EM. For convenience, we refer to both types of maps as “density maps” in this paper.

#### Conformational sampling for residues

qFit 3 exhaustively samples residue conformations in three stages: backbone sampling, Cα-Cß-Cγ bond angle sampling (for certain residues), and side-chain sampling (**Figure 5A)**. These are all enabled by default, but can be individually disabled via command-line options.

#### Backbone sampling

qFit 3 samples backbone conformations by means of a nullspace inverse kinematics algorithm ^28,35,43^. Backbone sampling for each residue extends to neighboring residues, two on each side. Backbone sampling is not performed if a residue lacks two neighbors on both sides (e.g., close to terminal residues). The Cß atom of the residue of interest (or O atom for Gly) is moved in the direction of the major and minor axes of its thermal ellipsoid. By default, three amplitudes for this sampling are used (0.1 + σ, 0.2 + σ, 0.3 + σ), where σ is randomly selected in the interval [-0.125, 0.125].

The amplitude scaling factor and the maximum value of σ can be defined at input. In total, three amplitudes times six directions = 18 positions for the Cß (O in case of glycine) are tested. The five-residue fragment is then deformed using nullspace inverse kinematics and dihedral angle degrees of freedom. The input conformation is also added to the ensemble, leading to 19 backbone conformations after backbone sampling. Peptide flips ^35^ are not yet implemented in qFit 3.

#### Cα-Cß-Cγ bond angle sampling

For amino acids with large planar aromatic groups (Phe, Tyr, Trp, His), qFit samples around the Cα-Cß-Cγ bond angle of the 19 backbone conformations resulting from the previous sampling step. For each conformation, we sample the Cα-Cß-Cγ bond angles as follows: [θ - 7.5°, θ - 3.75°, θ, θ + 3.75°, θ + 7.5°]. Both the range and the step of the bond angle sampling can be adjusted via command line. This step expands the number of sampled conformations to 95 for the large planar aromatic residues.

#### Side-chain sampling

Side-chain sampling in qFit 3 is performed by iteratively rotating around the χ angles of ideal rotamers. The protocol begins by rotating around χ_1_. For each of the (19 or 95) backbone conformations, we rotate around each of the rotamers for the target residue in the penultimate rotamer library ^25^. For each rotamer, we explore a sampling window using a rotamer neighborhood of [-60°, +60°] at 10° intervals. Both the sampling window and the step size can be defined via command-line options. For the default parameters, at most 19*5*(8+1)*13 = 11,115 conformations are generated (with either Phe, Tyr, Trp, or His), which provides a balance between performance and accuracy. From this set, we remove conformations that lack support from the subtracted density map (voxel with minimum density intensity < 0.3 e^-1^ Å^-3^), conformations that contain self-collisions (based on hard spheres), and conformations that are redundant (using an all-atom RMSD threshold of 0.01 Å). These exclusion strategies can be adjusted via command-line options. For protein and ligand atoms, B-factor sampling is also a non-default option.

Once the backbone and χ_1_ sampling is complete, the protocol initiates a selection step based on our optimization strategy (see Optimization protocol for more details). We select all atoms starting from the backbone up to the atoms involved in the χ angle being sampled (χ_1_ in this first iteration). The remaining atoms are rendered inactive, and their density contribution is not taken into account during optimization. Up to five conformers can be selected at each iteration, which then serve as the basis for sampling of subsequent χ angles.

From the second iteration onwards, we sample up to two χ angles simultaneously (also defined at command line). After sampling χ_i_ we exclude unsupported, clashing, and redundant conformers (as outlined above) and use this filtered conformer ensemble to sample around χ_i+1_. In the worst case scenario (Arg), χ_i_ leads to 5*(34+1)*13 = 2,275 conformers and up to 2,275*(34+1)*13 = 1,035,125 conformations are produced for χ_i+1_. In practice, this number of conformations is never produced owing to redundancy. We limit the number of conformations that can be used during optimization to 15,000 for computational efficiency and memory (RAM) constraints. If sampling two χ angles in a single iteration leads to more than 15,000 conformers, we reduce sampling to a single χ for that iteration. Side-chain sampling concludes when all χ angles have been sampled.

#### Conformational sampling for ligands

Ligand sampling in qFit 3 is performed in two steps: a local rigid body search followed by an iterative step which samples the degrees of freedom about the flexible areas of the ligand ^32^ (**Figure 5B)**. For the local search, we identify all possible roots, i.e. rigid fragments of atoms. Rigid fragments are defined as a set of connected atoms that do not contain a rotatable bond. We sample conformations starting from each possible root. Around the center of each ligand root, we test 100 possible rotations, by sampling rotation space in intervals of [0°, 10°]. For each rotation, we enumerate possible translations for x, y, and z coordinates in the interval [-0.2 Å, 0.2 Å] at 0.1 Å increments. The local search leads to 100(rotations)*125(translations) = 12,500 conformers. We then exclude conformers that do not have support from the density (voxel with minimum density intensity < 0.3 e^-1^ Å^-3^) and conformations that are redundant, using an all-atom RMSD threshold cutoff of 0.01 Å. Additionally, conformers with internal (ligand) or external clashes (receptor) are removed using a spatial hashing algorithm, which efficiently converts the 3D coordinates to a 1D hash table to determine if the sampled portion of the ligand occupies the same spatial coordinates as any other part of the ligand and/or receptor. After this exclusion step, remaining conformations are used as input for the optimization routine (see below), which selects up to five conformers of each root to best represent the local density.

Still treating each root independently, we take the root fragments selected by the local rigid body search and “expand” each fragment to the full ligand, by iteratively sampling around rotatable bonds. The protocol follows a rotatable bond hierarchy from the root to the extremities of the molecule. For each rotatable bond, we sample all angles in a [0°, 360°] interval at 10° increments. Two rotatable bonds are sampled at a time, leading to 5*36*36 = 6,480 conformations per iteration. At each iteration, we exclude conformers that do not have support from the density (voxel with minimum density intensity < 0.3 e^-1^ Å^-3^), those with an all-atom RMSD of <0.01 Å, or that contain internal or external clashes. After exclusion, qFit uses the optimization routine to select up to five conformers to be used for the next iteration. After all rotatable bonds have been sampled, up to five conformers can be output for each root. One final optimization step is used to select up to five consensus conformers from the pool of conformers produced across all roots.

#### Optimization Protocol

We frame the problem of selecting a subset of conformers that best represents local density as an optimization problem. Each conformer has an occupancy *ω*_*i*_ associated with it. The vector of all occupancies ***ω***^***T***^ contains the variables for the optimization, with the extra constraints that *ω*_*i*_ are non-negative and their sum lies in the unit interval. We optimize real-space residuals, calculated from the observed density (*ρ*^*obs*^) against the occupancy-weighted sum of the calculated densities (*ρ*_i_^calc^) for all conformers. We can formulate this problem as constrained quadratic optimization:

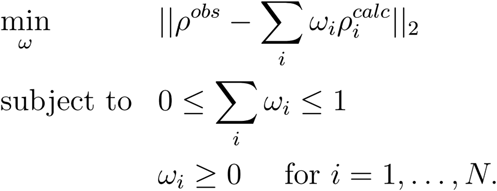

Residuals are calculated over all voxels within (0.5 + resolution / 3) Å from any active atoms across all input conformers. To prevent overfitting conformers with arbitrarily small occupancies, we require a threshold constraint on the occupancies, turning the problem into a mixed-integer quadratic program (MIQP):

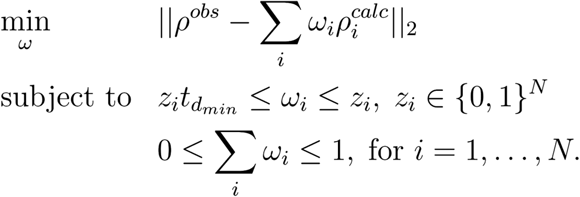

Note that this ensures that the number of conformers selected is at most 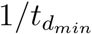. The optimal threshold parameter is determined using a penalized-likelihood criteria (see below). An MIQP is NP-hard, thus applying an MIQP solver directly to the conformers output from our sampling step is computationally inefficient ^28,35^. Applying a QP solver to the thousands of conformers output from our sampling routine, and then selecting the QP-fitted conformers with non-zero occupancy as input for MIQP, allows for near-optimal solutions to be calculated within a tractable time. Our protocol uses cvxopt (https://cvxopt.org/) and a proprietary, freely available implementation of the IBM ILOG CPLEX Optimization Studio (Python API, version 12.10) to solve QP and MIQP programs.

#### Achieving parsimony by means of the Bayesian Information Criterion (BIC)

To prevent overfitting and to ensure optimal model selection, we use the Bayesian Information Criterion to decide on model complexity. For every optimization call in qFit, we iteratively test increasing values of the threshold parameter 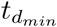 and determine if the gain of information justifies the use of a more complex model. We fit iteratively, allowing the maximum number of conformers to vary from 1 up to 5 conformers ranked according to real-space correlation. For each iteration, we use our combined QP/MIQP routine to optimize the real-space residual sum of squares (RSS). We calculate the BIC for each level of complexity according to the following formula:

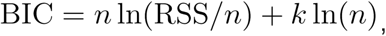

where *n* is the number of voxels in our resolution-dependent mask (see previous section for details) and 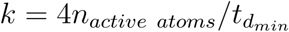 is the number of parameters in the model.

Each active atom has four parameters: x, y, z, and B-factor. Note that the occupancies are variables and not parameters. The factor 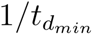 is a proxy for model complexity and imposes a limit on the maximum number of conformations. We select the number of conformers that minimizes the BIC.

#### Parallelization

qFit 3 can be run individually for a single residue or ligand of interest, or in parallel across a whole protein using Python’s *multiprocessing* module to spawn embarrassingly parallel subprocesses that run qFit across all residues in a target protein.

#### Validation metrics

For each residue/ligand modeled by qFit 3, we output several validation metrics, which include the BIC and the related Akaike information criterion AIC = 2*k* + *n* ln (RSS) with and k as above. qFit 3 also reports a confidence interval for the real-space cross-correlation of the proposed conformers. The confidence interval is calculated from the Fisher z-score of the real-space cross-correlation *r*^51^:

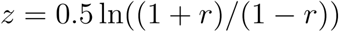

Note that the z-score is approximately normally distributed with a standard deviation of 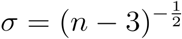, where *n* is the number of voxels in our resolution-dependent mask around the set of conformers being assessed. qFit 3 reports the 95% confidence interval *z* ± 1.96σ for the cross-correlation. Overlapping intervals suggest that the gain in cross-correlation is statistically not significant; we cannot reject the null hypothesis that the cross-correlations are the same at 95% confidence.

These auxiliary validation metrics are not used to filter results, but provide a guideline for balancing gain of information vs. model complexity.

#### Building an internally consistent structural model

In the procedure above, residues are modeled independently, i.e., without taking into account multiconformer models for neighboring residues. This leads to two modeling inconsistencies. First, consecutive residues may have different occupancies for each altloc, or even a different number of alternate conformations. Second, alternate conformers of (not necessarily consecutive) side chains in a spatial neighborhood can clash owing to inconsistent assignment of altloc identifiers. To resolve these two inconsistencies, we execute two routines: qFit-segment, which addresses the problem of inconsistency along the backbone, and qFit-relabel, which resolves clashing alternate conformers between neighboring residues by reassigning altloc labels.

The qFit-segment routine starts by identifying all segments along the backbone for which all residues have at least two backbone conformers. To mark the start and end points of such backbone segments, we identify residues for which either (a) a single conformer was output, or (b) where the backbone Cα and O atoms of that residue’s conformers do not deviate by more than 0.05 Å. A segment is then delimited by these single-backbone-conformer residues. To create consistent segments, we proceed iteratively. We break the segments in fragments of up to 4 residues (adjustable via the command line). We enumerate all possible combinations of conformers for the fragment, which at worst case leads to 5^4^ = 625 possible conformers. We use our optimization strategy (QP/MIQP iteratively, using the BIC) to select up to five conformers per fragment based on optimal fit to the experimental map (not based on covalent geometry). To ensure consistency with the PDB file format and compatibility with refinement software, we duplicate conformers for some residues within a fragment as needed to ensure that all consecutive residues have the same number of backbone conformers. Once all 4-residue fragments have been modeled in this fashion, we proceed to enumerate all possible combinations of such length-4 fragments. This leads to fragments of at most length 16, and, again, at worst case 5^4^ = 625 possible conformers. We continue to iterate in this fashion, enumerating all possible combinations and solving/modeling, until the segment is completed. The output of the qFit-segment routine is segments, each with up to five conformers, for which the backbone is consistent, i.e., for which all atoms for each conformer have the same label and occupancy.

Next, qFit-relabel relies on simulated annealing (SA) optimization of a Lennard-Jones potential to reassign altloc labels. We calculate the pairwise Lennard-Jones potential across every atom of all conformers output by qFit. Parameters for the Lennard-Jones calculation were taken from the Amber ff99SB forcefield ^52^. The procedure selects five segments at random (a segment can include a single residue in this case) and randomly shuffles their labels. We then assess the change in the Lennard-Jones potential and either accept or reject this move. The probability of accepting an unfavorable move is defined as:

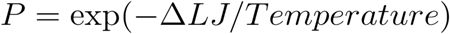

The temperature begins at 273 (arbitrary units), and is decreased by 10% every 10,000 perturbations. By default, 100,000 perturbations are sampled during relabeling. Benchmarking suggests that this value is sufficient for the scoring function to converge (data not shown). The output of the relabeling routine is a multiconformer model with up to five conformers per residue, in which backbones are consistent and in which alternate conformers for side chains are not clashing.

#### Final refinement script

We performed iterative refinement on the qFit multiconfomer models using version 1.18 of the Phenix software suite ^33^ to normalize the initially distorted covalent geometry, to ensure that the output models are properly fit into density (**Supplementary Figure 3)**, and to remove any unnecessary conformers.

For X-ray crystallography structures, this iterative refinement protocol uses the *phenix*.*refine* executable (script name: qfit_final_refine_xray.sh). The initial round of refinement is done without hydrogens and uses the strategy=*individual_sites. We then (re-)add hydrogens to the model ^53^. The next rounds of refinement use the following parameters: strategy=*individual_sites *individual_adp *occupancies, number_of_macro_cycles=5. At each iteration, we remove all conformers for which the occupancy fell below a cutoff of 0.09. This iterative cycle continues for as long as atoms are being removed due to this occupancy cutoff criterion. We then perform one last refinement round.

For cryo-EM structures, we use a similar refinement protocol as described above, but using *phenix*.*real_space_refine* ^36^ (script name: qfit_final_refine_cryoem.sh). All rounds of real-space refinement use the default parameters.

### High-performance and cloud computing

qFit is capable of scaling from single laptops to large high-performance computing clusters. The following instructions enable qFit on Amazon’s AWS, and should readily generalize to other cloud providers and RPM-based Linux distributions.

We describe configurations at two different scales: a single instance and an autoscaling cluster with a free master instance.

#### Single instance

Launch an instance that will be used to execute qFit. AWS’s `c5.9xlarge` instance has an appropriate number of cores and amount of memory for most proteins.

The following Bash script, reproduced from `docs/aws_deploy.sh` in the qFit repository, installs qFit and its dependencies within a conda environment:

~~~
#!/usr/bin/env bash
# Tested on Amazon Linux 2, but should work on most RPM-based Linux distros # install Anaconda RPM GPG keys
sudo rpm --import https://repo.anaconda.com/pkgs/misc/gpgkeys/anaconda.asc
# add Anaconda repository
cat <<EOF | sudo tee /etc/yum.repos.d/conda.repo [conda]
name=Conda baseurl=https://repo.anaconda.com/pkgs/misc/rpmrepo/conda enabled=1
gpgcheck=1 gpgkey=https://repo.anaconda.com/pkgs/misc/gpgkeys/anaconda.asc EOF
sudo yum -y install conda sudo yum -y install git gcc
source /opt/conda/etc/profile.d/conda.sh conda create -y --name qfit
conda activate qfit
conda install -y -c anaconda mkl
conda install -y -c anaconda -c ibmdecisionoptimization cvxopt cplex
git clone https://github.com/ExcitedStates/qfit-3.0.git cd qfit-3.0/
# Optionally, uncomment the following line to set a specific version of qFit #git checkout v3.2.0
pip install.
~~~

Consider creating an image of the instance at this point to avoid executing the above script each time an instance is launched from a base instance.

After installation, it is necessary to execute `source`

~~~
/opt/conda/etc/profile.d/conda.sh
~~~

to set up conda within your Bash shell then activate the conda environment by executing

~~~
conda activate qfit.
~~~

Using the example described it qFit’s `README.md`, alternative conformers for all residues in 3K0N can be calculated by executing

`qfit_protein 3K0N.mtz -l 2FOFCWT,PH2FOFCWT 3K0N.pdb -p 36`

for `3K0N.mtz` and `3K0N.pdb` in the current working directory, utilizing up to 36 cores.

#### Autoscaling cluster

Additionally, ParallelCluster can be used to create an autoscaling cluster to maximize efficiency of cloud resources. See Supplementary Methods for details.

## Acknowledgements

SAW is supported by NSF GRFP 16501133. JSF is supported by NIH GM123159, NIH GM124149, and NSF STC-1231306. DAK is supported by NIH R35GM133769.

