## Supplemental Text and Figures for "qFit 3: Protein and ligand multiconformer modeling for X-ray crystallographic and single-particle cryo-EM density maps"

### Supplementary Information

#### Supplementary Methods

##### High-performance and cloud computing

###### Autoscaling cluster

###### Cluster creation and configuration

ParallelCluster is a suite of officially supported open-source tools used to create an autoscaling cluster on AWS.

After installation, `pcluster configure` provides a setup assistant to configure a cluster. A series of prompts guides the user through selection of region, scheduler, operating system, minimum and maximum size, master and compute instance type, and network configuration. These instructions assume selection of Slurm as the scheduler and Amazon Linux 2 as the operating system.

The following `[cluster]` section of the configuration file (saved at `~/.parallelcluster/config` on Linux) represents reasonable settings:

```
[cluster default]
key_name = ###redacted###
base_os = alinux2
scheduler = slurm
master_instance_type = t2.micro
cluster_type = ondemand
compute_instance_type = c5.9xlarge
max_queue_size = 10
maintain_initial_size = false
vpc_settings = default
post_install =
https://raw.githubusercontent.com/ExcitedStates/qfit-3.0/master/docs/aws_deploy.sh
```

This cluster will always run a `t2.micro` master instance, the first 750 hours per month of which are free, and a variable number of `c5.9xlarge` compute instances. While the scheduler's queue is empty and all jobs have finished, no compute instance will be running; when a job is submitted, a new compute instance will be launched so long as the total number would not exceed `max_queue_size`. New instances will download and execute the file at `post_install` URL, installing and configuring qFit.

For reduced costs in exchange for risk of job termination, `cluster_type` can be set to `spot` instead of `ondemand`. Spot pricing and risk of interruption are variable and depend on instance type, which should be considered when selecting compute instance type.

The cluster can be created with the command `pcluster create default`, accessed via SSH with `pcluster ssh default` and deleted with `pcluster delete default`.

##### Job submission

A job performing the same computation as described for the single-instance configuration can be created with the following Bash script:

```
#!/bin/bash
#SBATCH --job-name=3K0N
#SBATCH --output=3K0N.out
#SBATCH --error=3K0N.err
#SBATCH --qos=normal
#SBATCH --ntasks=1
#SBATCH --cpus-per-task=36
#SBATCH --time=240:00

source /opt/conda/etc/profile.d/conda.sh
conda activate qfit
qfit_protein 3K0N.mtz -l 2FOFCWT,PH2FOFCWT 3K0N.pdb -p 36
```

Assuming the above is saved to `3K0N.job`, the job can be submitted by executing `sbatch 3K0N.job`.

If many jobs need to be submitted, we recommend creating one directory per protein, with each containing an MTZ and PDB file. If the following file is saved as `job_gen.sh` in the directory containing `3K0N/` (which contains `3K0N.mtz` and `3K0N.pdb`) and `4K0N/` (containing `4K0N.mtz` and `4K0N.pdb`), jobs for 3K0N and 4K0N can be created and submitted by executing `./job_gen.sh 3K0N 4K0N`.

```
#!/bin/bash

STARTDIR=$(pwd)

for protein in "$@"
do
	protein_dir=$STARTDIR/$protein
	job_file=$protein_dir/$protein.job
	echo $protein
	echo $protein_dir
	echo $job_file
	echo "#!/bin/bash
#SBATCH --job-name=${protein}
#SBATCH --output=${protein_dir}/${protein}.out
#SBATCH --error=${protein_dir}/${protein}.err
#SBATCH --qos=normal
#SBATCH --ntasks=1
#SBATCH --cpus-per-task=36
#SBATCH --time=240:00
```

```
source /opt/conda/etc/profile.d/conda.sh
conda activate qfit
qfit_protein ${protein_dir}/*.mtz -l 2FOFCWT,PH2FOFCWT ${protein_dir}/*.pdb -p 36
" > $job_file
	cd $protein_dir
	sbatch $job_file
	cd $STARTDIR
done
```

### Supplementary Figures

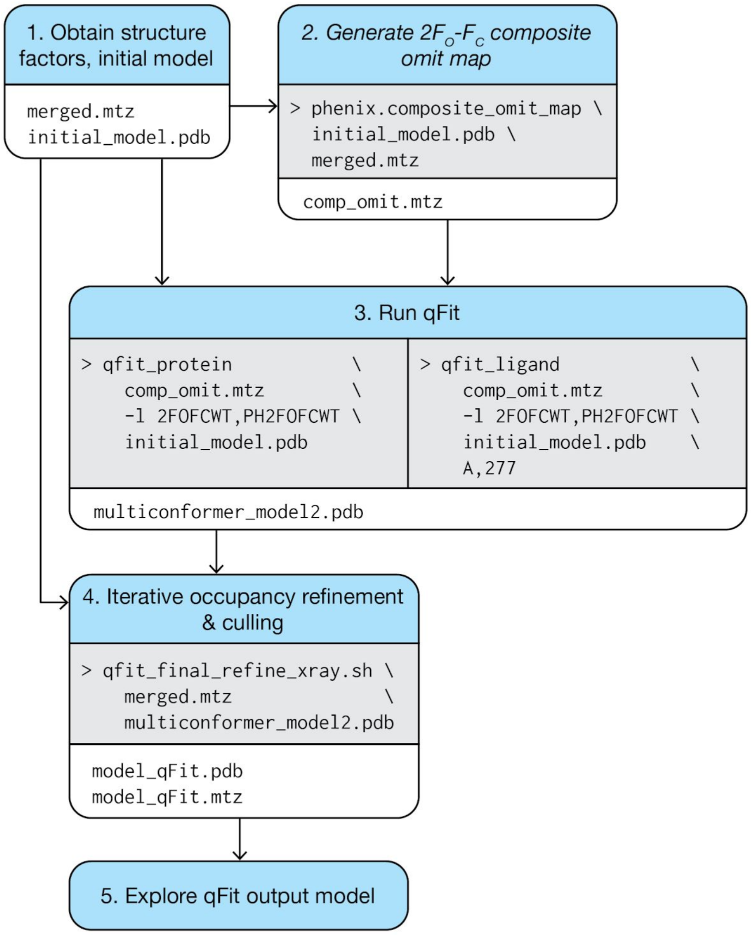


**Supplementary Figure 1:**
A flowchart for typical use of qFit with X-ray data.

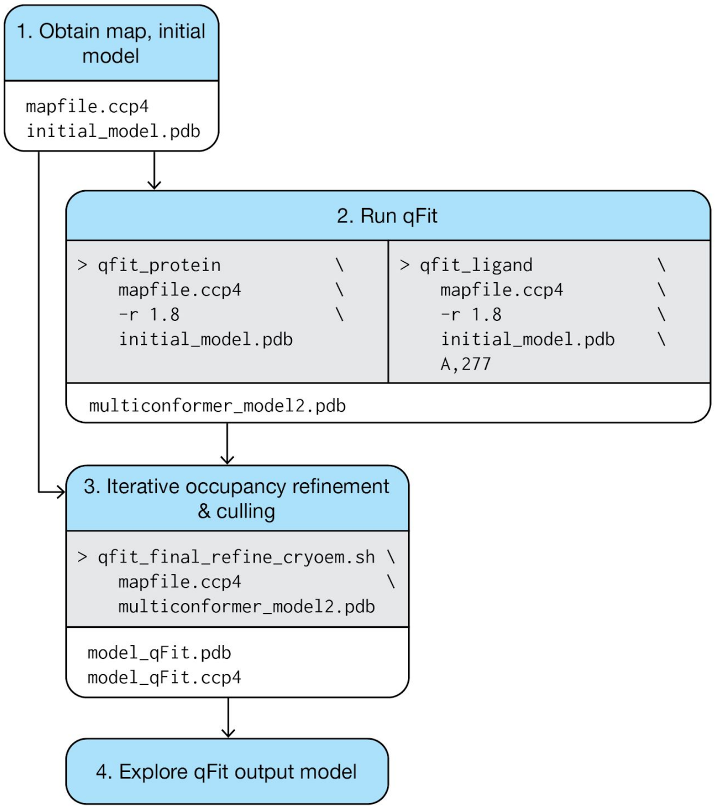


**Supplementary Figure 2:**
A flowchart for typical use of qFit with cryo-EM data.

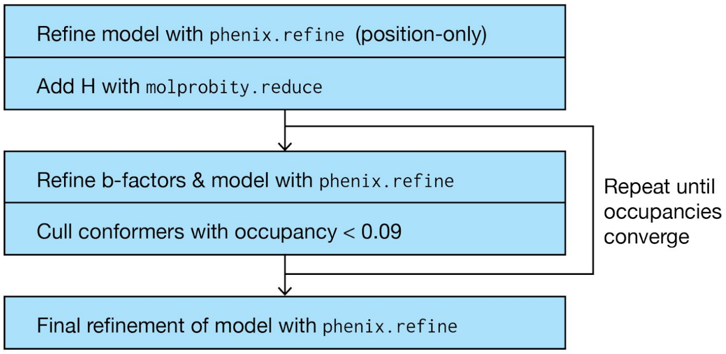


**Supplementary Figure 3:**
A flowchart for the recommended final refinement procedure. This was used for all structures modeled by qFit in this paper, and is contained in both `qfit_final_refine_xray.sh` and `qfit_final_refine_cryoem.sh`.
